# Geometry-preserving Expansion Microscopy microplates enable high fidelity nanoscale distortion mapping

**DOI:** 10.1101/2023.02.20.529230

**Authors:** Rajpinder S. Seehra, Benjamin H.K. Allouis, Thomas M.D. Sheard, Michael E Spencer, Tayla Shakespeare, Ashley Cadby, Izzy Jayasinghe

**Affiliations:** School of Biosciences, Faculty of Science, The University of Sheffield, Sheffield S10 2TN, UK; Department of Physics and Astronomy, The University of Sheffield, S10 3RH, UK

**Keywords:** expansion microscopy, correlative microscopy, super-resolution, 3D printing, distortion mapping

## Abstract

Expansion microscopy (ExM) is a versatile super-resolution microscopy pipeline, leveraging nanoscale biomolecular cross- linking and osmotically driven swelling of hydrogels. In its current implementation, ExM remains a laborious and skill-intensive technique, involving manual handling of the hydrogels that can compromise the integrity of the gel matrix and diminish reproducibility. The lack of protocols to constrain the gel orientation during this process lends to challenges in tracking gel isotropy during or after the swelling. We have developed a bespoke microplate system capable of carrying out the entire ExM workflow within each well. The microplates enable *in situ* image acquisition and eliminate the need for direct physical handling of the hydrogels. The preservation of the gel geometry and orientation by the design of the microplate wells also enables convenient tracking of gel expansion, pre- and post-ExM image acquisition, and distortion mapping of every cell or region of interest. We demonstrate the utility of this approach with both single-colour and multiplexed ExM of HeLa cells cultured within the microplate wells to reveal nuclear and sub-plasmalemmal regions as distortion-prone structures.

## Introduction

Super-resolution microscopy methods have considerably shifted the capabilities of optical bioimaging, from cellular or sub-cellular scales to the true molecular scale. Whilst the conventional stochastic (e.g., STORM, PALM, and PAINT) and deterministic (e.g. 4Pi, STED, and SIM) approaches to super-resolution have pioneered many of the breakthroughs in this technology, access to the specialist optical instruments and mastery of the underpinning photochemistry continue to limit its uptake. Expansion microscopy (ExM) has offered a radically different strategy to achieving super-resolution [1]. By combining molecular cross-linking chemistries with osmotically swellable polyacrylamide hydrogels, ExM enables the *physical* magnification of the ultrastructure. Super resolution can therefore be achieved through fluorescence imaging of an inflated facsimile of the structure with relatively conventional (e.g., confocal) imaging systems.

The utility of ExM has grown with the development of a range of ExM recipes, each with improvements in either (i) accessibility of key reagents and components [2], (ii) expansion factor towards greater resolution [3, 4], (iii) isotropy of gel expansion towards higher fidelity sample expansion [4-6], and/or (iv) compatibility with different sample species [7], biomolecules [8], constituents [9], and sample formats [10]. Combining of ExM chemistries with established super-resolution modalities has given rise to ExM variants such as ExSTED [11], ExSIM [12], ExSMLM/Ex*d*STORM [13], EExM [14, 15], and one-nanometre expansion (ONE) microscopy [16], allowing users to compound the resolution gain between the optical, computational and hydrogel enhancements. Finally, the rapidly growing range of molecular probes and staining protocols [9, 17], particularly the adaptation of nondescript counter stains [18, 19], have also made ExM a highly versatile route to super-resolution microscopy.

Despite the variations of ExM, the workflow of sample preparation, particularly from the gelation to image acquisition stages, remain highly reliant on manual handling of the gel and user skill [20]. Of note, the method involves repeated manual handling of the gel during the transfer, osmotic swelling, and trimming of the gel blocks. Skill is required to minimise distortion, damage, and/or tearing of the hydrogels and the structures imprinted within. Further to this, spatial heterogeneities in the gel matrix, proteolytic digestion, and intrinsic stiffness within the sample are known to give rise to anisotropies in the gel expansion [15, 21]. The same phenomena can also give rise to sample-to-sample and region-to-region variations in expansion factor, usually measured by changes in the overall gel dimensions [14]. Additional strategies such as, embedding DNA-origami calibrants [6], using biological structures and shapes intrinsic to the samples (e.g. Nuclear pore complexes [22] or muscle sarcomeres [14]), and using patterned fluorescence photobleaching [23] or gridded substrates that impart trackable imprints on the gel [24]. Nevertheless, imaging the same regions or structure at the pre- and post-expansion states [25] remain the Gold standard for assessing expansion factor and any distortions. Due to the manual nature of the gelation and gel expansion steps, sequential pre- and post-ExM imaging currently remains a highly skilled and time-consuming mode of ad-hoc validation.

Array-based ExM promises to remove or reduce the manual handling of the hydrogels and add scalability to this method which, at present, remains a low-throughput imaging modality. In this study, we present the adoption of 3D printing and rapid prototyping as a strategy to develop a bespoke microplate-based ExM protocol that also preserves hydrogel geometry and orientation to enable trackable, pre- and post-ExM imaging of every sample and region of interest.

## Results and Discussion

Expansion Microscopy is achieved through a series of chemical anchoring, gelation, digestion, and expansion steps after a sample has undergone fluorescence or immunofluorescence staining (Fig 1A). In its typical implementation, coverslips containing cell or tissue samples are manually lowered onto an aqueous mix of gel monomers, polymerization catalysts and inhibitors. The polymerized gel may be transferred to a chamber or plate for the proteolytic digestion and washes, and then transferred again to a large petri dish for the osmotic swelling in the expansion stage. Expanded gels are trimmed to fit the size of an imaging chamber before being placed on the microscopes. The latter three steps each involves manual handling and transfer of the gel. Considerable skill and experience are required to ensure that the integrity and isotropy of the sample are preserved.

**Figure 1.**
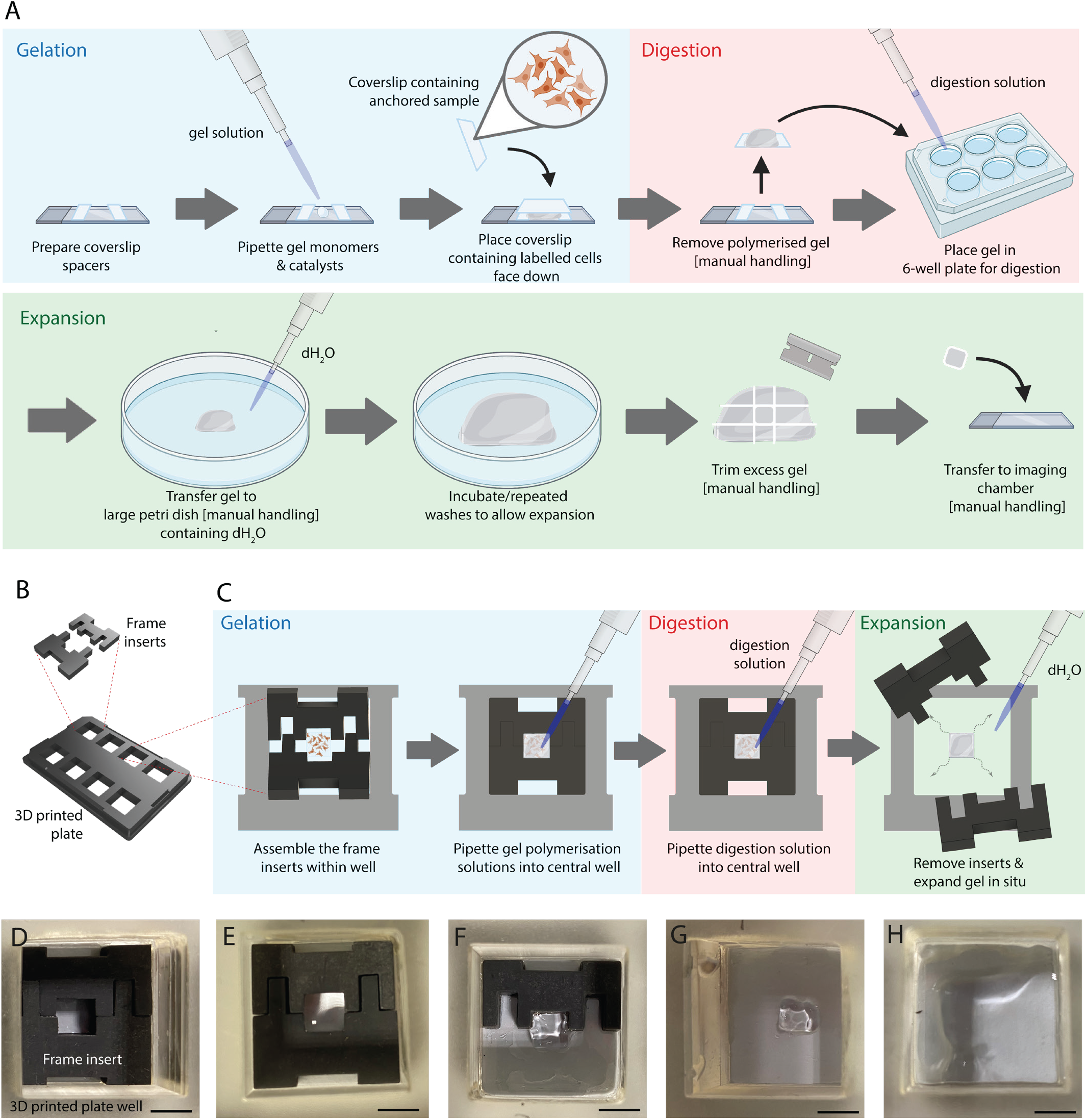
Conventional and microplate-based implementation of ExM. **A**. The conventional ExM protocol involves the three main stages of ExM, Gelation, Digestion, and Expansion. Manual handling and/or transfer of the gel is likely in the post-gelation, post-digestion and post-expansion stages. **B**. The design of 3D printable microplate, consisting of eight wells, and a pair of silicone frame inserts which assembles into each well. **C**. In the microplate-based ExM workflow, the frame inserts are pushed to the bottom of the well either prior to seeding the cells in the central region of the well or following immunostaining. The gel polymerisation and digestion steps take place within the central well. The frame inserts are removed to release the square shaped gel prior to the Expansion step. Carefully adding deionised water (dH_2_O) allows the gel to expand 4x to fill the main well. Shown are a series of photographs of: **D**. the assembly of inserts in the well, **E**. The forming of the gel within the central gel casting well, **F**. the releasing of the gel from the frame inserts, **G**. the released gel (pre-expansion), and H. the expanded gel filling the overall well. Note, excess liquid has been removed from the wells for clarity of the photographs. Scale bars: 5 mm.

We developed a 3D-printable, 8-well microplate and a set of silicone frame inserts to perform ExM within each well without having to remove the gel or the sample at any stage of the protocol (Fig 1B; the design model is included in the data supplement). Each well was a 2×2 cm square space and a No.1.5 glass coverslip bottom to allow direct imaging from underneath. Cells could either be cultured within the central portion of each well or the inserts placed after the cells were cultured and fixed *in situ* (Fig 1C). In the latter approach, it was important to blot away any residual liquid from the edges of the well to allow the silicone frame inserts to form a dry seal when pressed into the well. Similar to a jigsaw, each pair of frame inserts was assembled and seated within the chamber to form a vacant central region either before adherent cells were seeded or following the staining and anchoring steps. The central well (a 0.5 × 0.5 cm square) formed by the frame acted as the casting well for the hydrogel. The tight-fitting silicone frame inserts were pressed gently against the coverslip to form a dry seal. Once polymerized, the digestion solution and subsequent wash buffers could be pipetted into the same well. The frame was dismantled by leveraging tweezers gently against the slots (at the top and bottom of the frame) and removed from the well. Provided that the frame inserts formed a substantial seal against the dry portions of the coverslip, this step should release a square gel block at the center of the well; if not, careful scoring of the gel along the edge of the central well with a sharp scalpel will be necessary prior to the release of the gel block. To allow the square gel to expand, deionized water (dH_2_O) was gently pipetted into the same well taking care to prevent the gel from rotating. The protein-retention ExM [2] formula of the gel allowed it to expand to ∼ 4 times to fill the square space defined by the well. Fig 1D-1H is a sequence of photographs of the steps in performing ExM on HeLa cells cultured and fixed within a chamber. A 3D printable lid (included in data supplement) was also designed; however, lids of standard 98- or 6- well microplates were also compatible with our bespoke microplate.

The square geometry into which the gel is cast, along with the square shape of each well allowed us to maintain the expanded gel in its original orientation. Therefore, a cell or structure of interest originating at the top left corner of the casting well could be tracked reliably to the top left corner of the overall well following expansion (red asterisks in Fig 2A and 2B). The footprint of the microplate base was designed to match the industry-standard microplates, which allowed us to conveniently seat the plate within a standard stage plate of an inverted Zeiss Airyscan LSM 880 (Fig 2C). The No.1.5 glass coverslip bottom of each well permitted us to focus light through high numerical aperture objective lenses directly into the samples, *in situ*, both in the pre- and post-expansion image acquisition. Pre- and post-expansion Airyscan image pairs of HeLa cells stained directly with Alexa Fluor 488 NHS ester (Fig 2D&E) demonstrated the ability to conveniently track the same regions of the sample with minimal rotation.

**Figure 2.**
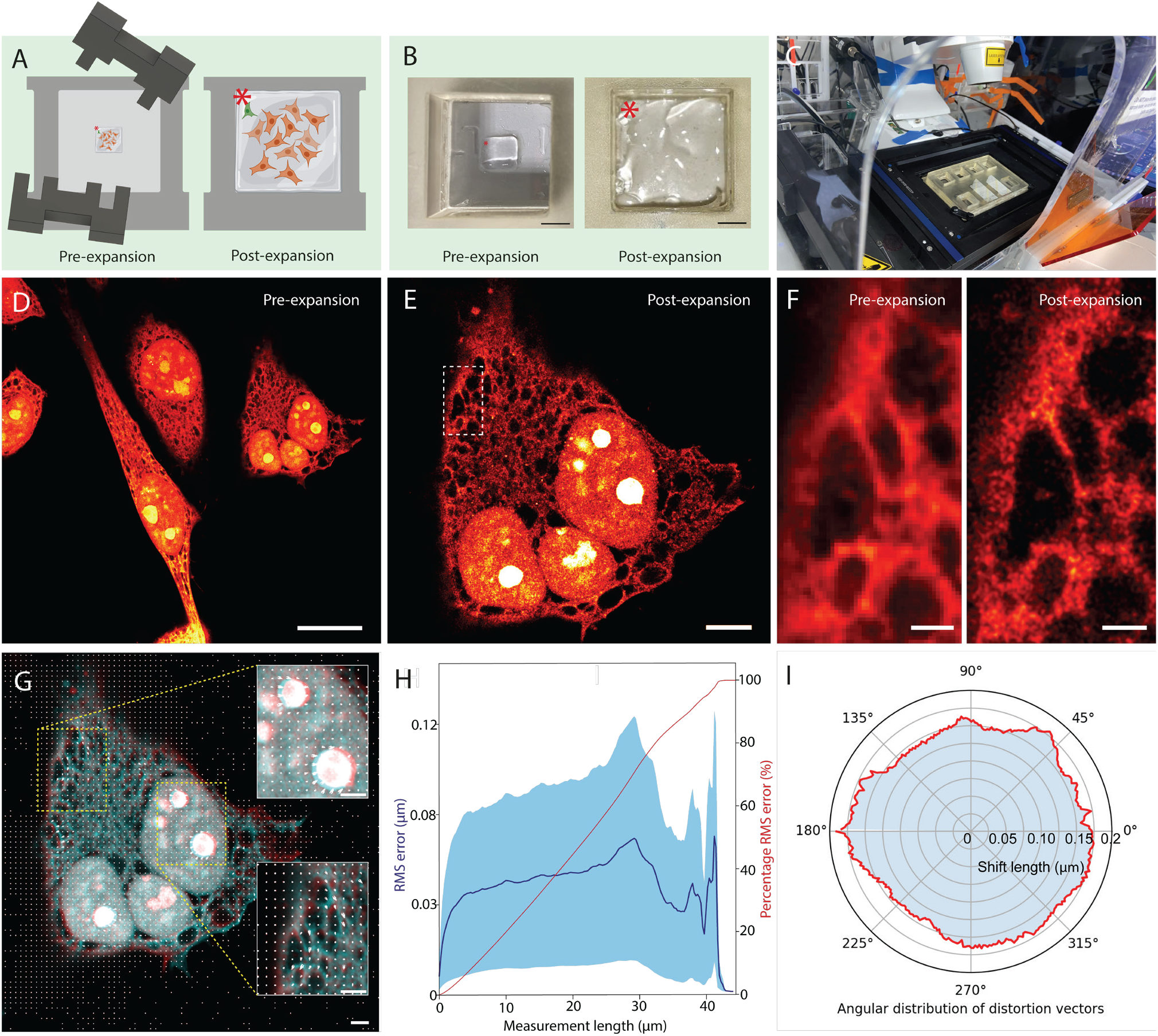
*In-situ*, pre- and post-ExM imaging and sub-cellular distortion assessment. **A**. Schematic diagram, and **B**. photographic illustrations of the square shape of the gel released from the frame inserts (left) and the gel following 4x in situ expansion filling the well (right). The preservation of the geometry and orientation of the gel means that the upper right corner of the sample within the casting well moves unambiguously to the upper right corner of the overall well (red asterisk). **C**. The coverslip bottoms of each well and the standardised base of the microplate allows it to be seated directly on a research grade Airyscan microscope for imaging into the gels at pre- and post-ExM stages. **D**. A pre-ExM and **E**. post-ExM Airyscan images of the same sample of HeLa cells stained with an AZ488 NHS ester. **F**. Magnified and re- scaled view of the region shown within the dashed lines in panel E at pre-ExM (left) and post-ExM (right) stages. **G**. Aligned and scaled overlay of the pre-ExM (red) and post-ExM (cyan) Airyscan images of the cell. The insets show magnified regions within the nucleoplasm and near cell periphery, marked by the dashed lines, featuring longer distortion vectors representing local distortions. **H**. Plots of RMS error (solid blue line; SD from distortion vectors shown in panel G) and cumulative normalised RMS error (red line) as a function of the measurement length scale. **I**. Polar plot of the average distortion vectors (radial axis) against directionality (angular axis). Scale bars: B: 5 mm, D: 20 μm, E: 5 μm, F: 1 μm and G: 2 μm.

This approach of pre- and post-ExM re-imaging enabled two principal modes of data visualization. Firstly, it was straightforward to *directly* compare the super-resolution image with the diffraction-limited standard to demonstrate the resolution improvement (Fig 2F) – as is often *not* shown in ExM studies. Secondly, cells or regions of interest could be directly scaled, registered and subject to a two-dimensional (2D) spatial distortion analysis. Fig 2G illustrates an overlay of the pre-expansion (red) and post-expansion (cyan) images, along with a 2D vector map of the local registration error (white arrows). This analysis allowed us to recognize the boundary of the cell, particularly near the interfaces of internal compartments such as the endoplasmic reticulum (ER) and the plasmalemma of the cell, and the intra-nuclear structures as the sub-cellular regions most prone to anisotropic expansion (see magnified view in insets of Fig 2G). Additionally, convenient pre- and post-expansion imaging allowed us to carry out a root mean square (RMS) error analysis (Fig 2H) to identify the length scales that feature the greatest spatial errors resulting from anisotropies or distortion on a cell-by-cell basis. It also allowed us to plot the distortion vectors in a polar plot that reports any systematic asymmetries or directionalities in the distortions present in the region of interest. The polar plot in Fig 2I confirmed that the directionality of distortion vectors largely followed a random uniform distribution.

The microplate based ExM approach further lent to multiplexed staining and imaging. Fig 3A&B and Fig 3C&D show two distinct dual-colour ExM experiments carried out within microplate wells. In the first, fixed HeLa cells were stained with a nondescript AZ488 NHS ester (upper row of Fig 3A) and co-immunostained for KDEL peptide sequence found in distal ER (lower row). Shown, are the pre-ExM (Fig 3A-i), scaled and aligned post- 4x ExM (ii), and overlay Airyscan images (iii) corresponding to the two labels. In addition to the visual overlays, wew could examine the distortion vector maps for each imaging channel separately, accounting for any distortions arising from linkage errors unique to each label. It also allowed us to recover distortion information in regions of the cell (in this example, the nucleoplasm and perinuclear regions) with the AZ488 NHS stain that were otherwise devoid of KDEL labelling. The comparison of the RMS error plots of the two channels showed that with more widespread stains such as the AZ488 NHS ester, we can report errors in longer length scales by comparison to relatively segregated or localized targets like KDEL (Fig 3A-v). A split-view comparison of the pre- and post-ExM images is shown in Fig 3B to illustrate the improvement in spatial resolution and contrast achieved *within* the sample of interest.

**Figure 3.**
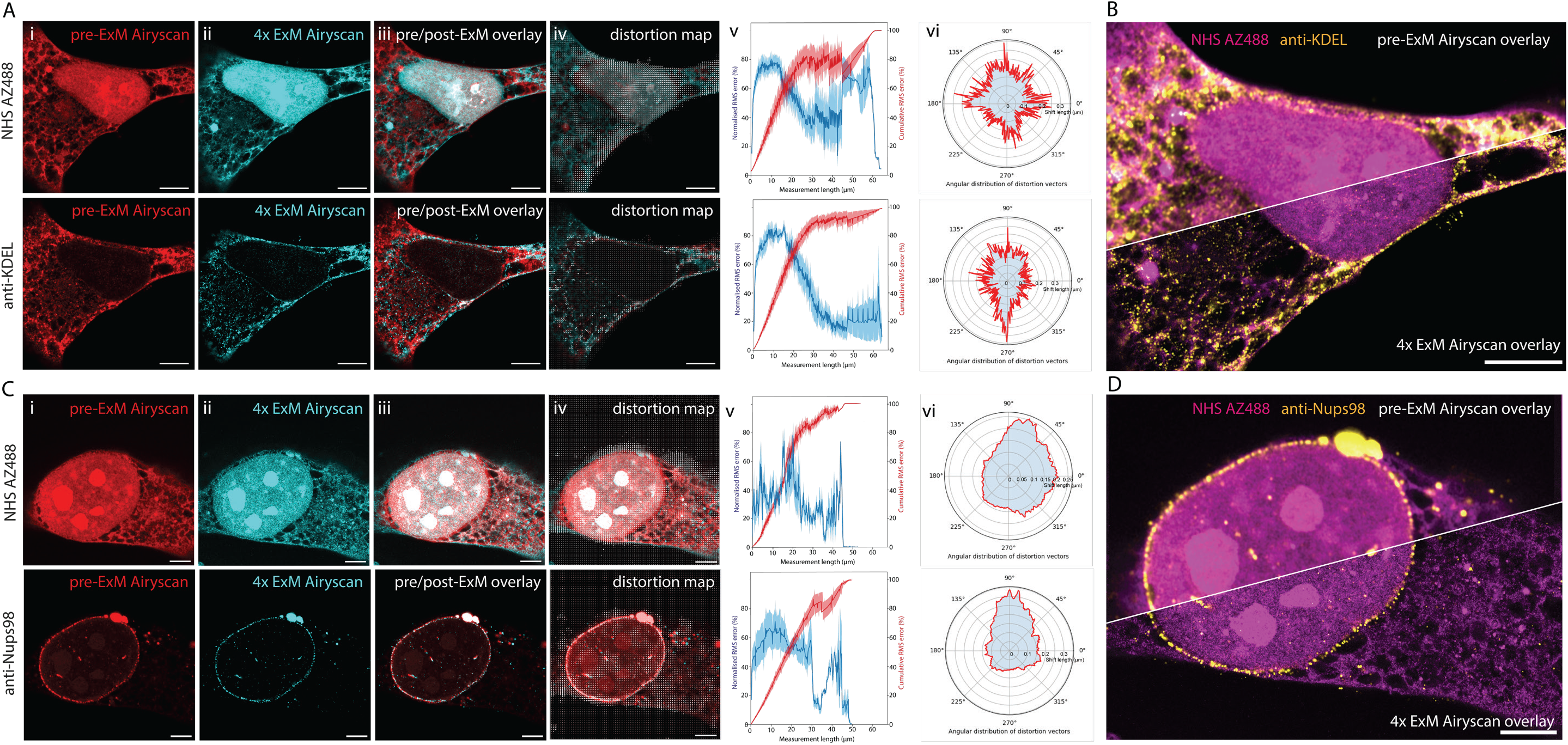
*In-situ*, pre-ExM and post-ExM multi-channel imaging of HeLa cells and local distortion assessment. **A**. A HeLa cell dual stained with AZ488 NHS ester (upper row) and anti-KDEL primary antibody (lower row). Shown, are **(i)** pre-ExM Airyscan, **(ii)** post-ExM Airyscan, **(iii)** Overlay of pre- and post-ExM images (pre-ExM in red and post-ExM in cyan), **(iv)** distortion map for each channel, **(v)** RMS error (blue; averaged between 8 image pairs from 8 cells across 4 wells from two plates; light shading indicates SD), and normalised cumulative RMS error (red) plotted against measurement length scale, **(vi)** polar plots of the average distortion vectors (radial axis) against directionality (angular axis). **B**. Split view of the overlay of the AZ488 NHS ester (magenta) and anti-KDEL (yellow) imaged with Airyscan at pre-ExM and post-ExM stages. RMS error plots are averaged between 6 image pairs from 6 cells across 4 wells from 3 plates. **C**. Equivalent views of a HeLa cell stained with AZ488 NHS ester (upper row) and a anti-Nups98 primary antibody (lower). **D**. Split view of the overlay of AZ488 NHS ester (magenta) and anti- Nups98 (yellow) imaged with Airyscan at pre-ExM and post-ExM stages. Scale bars: 5 μm

From the two-channel, pre- and post-ExM images, we consistently observed larger distortions both within and at the boundary of the nuclei – a feature that to our knowledge has not been reported in previous ExM studies. It is well known that the shape and volume of intact nuclei follow a non-linear relationship due to the restriction of solutes, particularly nucleic acid polymers across the nuclear envelope [26]. It was likely that restricted access for the gel monomers across the nuclear envelope was causing distortions in the nucleoplasm or nuclear envelope. We therefore performed the same sequence of analyses as above on HeLa cells co-stained with AZ488 NHS ester and Nup98, a key component of the nuclear pore-complex residing in the nuclear envelope (Fig 3C). The distortion maps for both channels (Fig 3C-iv - upper and lower) reveal prominent distortions both in the nucleoplasm and at the nuclear envelope, chiefly in the upward direction (see polar plots, Fig 3C-vi). We illustrate the visualization of pre- and post- 4x ExM Airyscan images (Fig 3D) with the recommendation that ExM image data must not only display the resolution improvement, but also accompanying channel-specific distortion vector maps as a part of responsible scientific practice.

As a rapidly evolving discipline of innovation, ExM continues to improve in its versatility and compatibility with various sample types. A significant domain of ongoing research and development is in leveraging ExM for high-resolution medical/biopsy imaging [27] as well as imaging complex, multicellular organisms [19] and their organs or tissues [28]. Whilst the microplate approach can be viewed as a gateway to improving through-put, our experience is that the fragility of polyacrylamide gels requires a level of care and attention that may be prohibitive for large sample numbers. The complexities of imaging with high numerical aperture objective lenses into non-refractive index matched samples such as these continue to require lengthy image acquisition sessions. However, we find the microplates are extremely useful for containing multiple samples within one array, for minimizing manual handling of the gels, and for enabling pre- and post-ExM imaging. We therefore conclude that its most valuable utility is its capacity for improving reproducibility.

## Materials and Methods

### Microplates and frame inserts

Both the microplate and the inserts were designed using Autodesk Fusion360 computer-aided design (CAD) software programme and exported as a .stl file. The microplate .stl CAD file was sliced using Chitubox Basic V1.9.0 slicer software and fabricated using an Elegoo Saturn 2 SLA 3D printer using the Translucent, low-odour translucent photopolymer resin (Elegoo) at 50 μm layer thickness, 5 bottom layers, 3.0 s standard exposure times, 20 s bottom layer exposure time, and no rest-time between steps. We have also reproduced qualitatively similar results with printing the same CAD file with Snapmaker 2.0 3D printer, sliced with Cura v5.0 (Ultimaker), standard black acrylonitrile butadiene styrene filament (ABS; Protopasta Ltd) at 0.24mm layer height, 18% infill density, 2 mm wall thickness, printing temperature of 245°C and bed temperature of 80°C. No.1.5 coverslips, sized 22 × 22 mm, (Menzel-Glaser, Germany) were adhered to the bottom using epoxy glass glue (Araldite) and cured under a UV hood. Frame inserts were laser cut from black, autoclavable silicone sheet rubber with a 3 mm thickness (custom ordered via Laser Web Ltd).

### Cell culture

HeLa cells were cultured in DMEM with high glucose, L-glutamine, HEPES, Phenol Red (Fisher Scientific Ltd, UK) and 10% FCS (v/v; LabTech International Ltd, UK), supplemented with 1% Penicillin-Streptomycin (v/v; containing 10,000 units/mL of penicillin 10,000μg/mL of streptomycin in a 10 mM citrate buffer; Fisher Scientific Ltd). Cells were kept at 37°C and 5% CO_2_ as standard and were used 2 days after passage.

Prior to cell seeding, the coverslip bottom of each well (pre-attached to the ExM plate) was coated with 0.01 mg/ml Poly-D-Lysine (code: 3439-100-01, Cultrex) for 2hrs at 37°C and 5% CO2. After removing the excess, cells were detached via 0.05% Trypsin-EDTA (Gibco, Fisher Scientific Ltd) and then plated at a density of 75,000/ml (1ml of the passaged mixture). Cells were kept in culture at 37°C and 5% CO2, with daily medium replacement till use after 2 days.

### Immunostaining

Fixation of HeLa cells was performed 2 days after plating, in 2% PFA (1ml fresh medium and 1ml of 4% PFA for final concentration of 2%) for 10 mins. Followed by 3 × 10 minute washes in fresh PBS. Cells were stored in PBS containing 0.5% bovine serum albumin (w/v) and 0.1% sodium azide (w/v) at 4°C till immunostaining steps.

Cells were permeabilized in PBS (137 mM NaCl, 2.7 mM KCl, 10 mM Na_2_KPO_4_, 2 mM KH_2_PO_4_, pH 7.3) + 0.1% Triton X-100, and subsequently blocked in PBS, 10% NGS (v/v) and 0.05% Triton X-100 (v/v).

Immunostaining was carried out in antibody incubation solution, with the addition of 200 μl primary antibody added to each well before incubation at 4°C overnight. The samples were then washed with fresh PBS for 3 times in 5 min steps, prior to secondary antibody application in 200μl for 2 hrs at room temperature. The samples were then washed in fresh PBS 3 times in 20 min steps.

For NHS ester application, after permeabilising the cells, 200 μl of 1:1000 NHS ester in ester staining solution (100 mM NaHCO_3_ + 1 M NaCl, pH 6, made up to 100 ml with dH_2_O) was added per well for 1 hr 30 mins at room temperature. Samples were washed 3 times with PBS in 20-minute steps prior to the blocking step.

### Antibodies used and catalogue number

#### Anchoring

Fluorophores were prepared for linking into the gel by incubating them at 4°C overnight in 250 μl of PBS and 1% Acryloyl-X (w/v; Catalogue number A20770, Fisher Scientific).

**Table 1:** Antibodies and fluorescent dye ester probes used for sample labelling

| Esters | Cat No. | Supplier | Dilution |
| --- | --- | --- | --- |
| NHS AZ488 | 1013-1 | Fluoroprobes | 1:1000 |
| NHS Alexa488 | a20000 | ThermoFisher Scientific | 1:1000 |
| **Primary Antibodies** |  |  |  |
| Anti-KDEL (Rabbit) | PA1-013 | ThermoFisher Sci | 1:500 |
| Anti-Nup98 (Mouse) | sc-74578 | Santa Cruz | 1:200 |
| **Secondary Antibodies** |  |  |  |
| Alexa 594 goat anti-rabbit | A11012 | ThermoFisher Sci | 1: 200 |
| Alexa 594 goat anti-mouse | A11005 | ThermoFisher Sci | 1:200 |

#### ExM Preparation

Cells in each well were washed with PBS for 3 times in 20-minute steps. They were incubated in 200μl of monomer solution (sodium acrylate, acrylamide, MBAA, NaCl, PBS, dH20) for 30 minutes at 4°C. The monomer solution was removed, and the Frame inserts were assembled in to each well, ensuring they are in good contact with the coverslip. 100 μl of 4X gel solution (containing 297 μl of monomer solution, 6 μl 10% TMED, 6μl 10% APS, 6μl PBS) was added to the centre of each Frame. The 10% APS and 10% TMED components were added to the gel mix immediately prior to dispensing into the wells to minimize pre-polymerisation of the gel before monomers were delivered into the cells. The plate was covered with either a bespoke lid or the lid of a 6-well plate allowing time for gel polymerisation. The plate was incubated at 37°C for 2 hrs.

#### Digestion and Expansion

The Frame inserts were gently removed from the wells with the aid of a tweezer, being careful in rare cases of gel adhering to the Frames. 1 ml of digestion buffer (containing 50 mM Tris, 1 mM EDTA, 0.5% Triton + 0.8 M guanidine HCl, made up to 100 ml with dH_2_O) along with 1% proteinase K (w/v; Sigma-Aldrich) was then added to each well and incubated on a rocker overnight at room temperature. Digestion should be performed for at least 12 hrs, with shorter times known to lead to gel damage. Each digested sample had an excess of dH_2_O to allow expansion whilst fully immersed. The dH_2_O was replaced at least 3 times, with 1 hour for each stage of expansion until approximately ∼ 4-fold expansion was achieved, i.e. until the gel reached the edges of the well.

#### Imaging

All images were obtained on a Zeiss LSM880 AiryScan (Carl Zeiss, Jena), using 10x 0.3 NA air Plan Apochromat objective and 40x 1.3 NA oil-immersion Plan Apochromat objective (both from Carl Zeiss, Jena). Imaging was performed in Airyscan mode with the gain and laser power adjusted for each sample to accommodate the fluorescence reduction occurring due to the spatial separation of fluorophores during expansion. Fluorophores were excited using Argon 488 nm and DPSS 561 nm laser lines. Selection of emission bands was performed through the spectral detector and recorded with the in-built GaAsP detector. Image acquisition was done through the associated Zen software and passed through the Airyscan post-hoc processing to obtain the final image.

#### Image analysis

Three avenues of image analysis were performed: pre-post expansion image alignment using ImageJ, distortion mapping using adjusted Python code from Truckenbrodt et al[25] (available via https://github.com/sommerc/expansion_factor_bftm), and Root-Mean-Square error (RMSE) data acquisition and plotting through a custom-written Python code. See supplementary movie 1 for a screencast of this analysis pipeline.

#### Pre-ExM and post-ExM image alignment

Pre-ExM and post-ExM images were aligned through the ImageJ plugin, “Linear Stack Alignment with SIFT”, with the transformation matrix visible. The intrinsic expansion factor (specific to the region of interest) was one of the outputs generated from the transformation matrix.

#### Distortion Analysis

The pre-ExM image was upscaled based on the estimated intrinsic expansion factor. Using the aligned image set, we determined the relative shift in coordinates of features present in the upscaled pre-ExM and the post-ExM images. Gunnar Farnebäck optical flow algorithm [29] was used to assess the relative shift between these images, as detailed previously by Truckenbrodt et al [25]. The output from this was used to produce distortion maps for the region of interest.

Using the aligned dataset, Otsu thresholding was performed on the post expansion image to generate binary data highlighting feature coordinates in the structure. The respective theoretical post alignment coordinates were then produced from the optical flow output respective to the feature coordinates. This series of theoretical “expanded”, pre-expansion coordinates, and actual post expansion coordinates were grouped by nearest neighbour distance. Nearest neighbour pairs between the coordinate sets were subtracted to obtain the associated difference between the length measurements. The original length between the coordinate points and the difference value were necessary to obtain the RMSE. Subsequently, the data was binned iteratively and the RMSE calculated for each difference value set associated with a bin position. In each iteration, the binning was adjusted to a pixel size that represented the length scale the RMSE was recorded against. In RMSE plots shown in Fig 3, multiple RMSE plots were averaged and shown with standard deviation recorded for each length scale.

The aligned images were commonly of size ∼ 2550×2550 pixels, hence the sequence of Otsu thresholding, erosion, closing, and the subsequent skeletonization of the binary image helped to reduce the 6 million+ coordinates down to between 30,000 - 200,000. The skeletonization also allowed for RMSE calculations to be constrained to regions that recorded a detectable pixel value above the typical background (i.e. regions with positive labelling).

## Code and data availability

The Python code, along with example datasets, are included in the data supplement. Also included in the data supplement are the .stl versions of the microplate and the frame inserts.

### Author Contributions

IJ secured the funding. IJ and RSS conceived and designed the experiments and tools; they also performed the manufacturing of the tools. RSS and BHKA performed the experiments, TS and MES provided the material. TMDS, AC, and IJ provided supervision. RSS and IJ performed the data analysis. IJ and RSS wrote the manuscript. All authors reviewed the manuscript.

## Acknowledgments

The biological samples demonstrating the methodology include the HeLa cell line. Henrietta Lacks, and the HeLa cell line that was established from her tumour cells without her knowledge or consent in 1951, have made significant contributions to scientific progress and advances in human health. The authors are grateful to Henrietta Lacks, now deceased, and to her surviving family members for their contributions to biomedical research.

The authors acknowledge funding from the UK Research and Innovation (MR/S03241X/1). Imaging work was performed at the Wolfson Light Microscopy Facility at the University of Sheffield. HeLa cell lines were donated by the Department of Infection, Immunity & Cardiovascular Disease of The University of Sheffield.

## References

1. Chen, F., P.W. Tillberg, and E.S. Boyden, Optical imaging. Expansion microscopy. Science, 2015. 347(6221): p. 543–8.

2. Tillberg, P.W., et al., Protein-retention expansion microscopy of cells and tissues labeled using standard fluorescent proteins and antibodies. Nat Biotechnol, 2016. 34(9): p. 987–92.

3. Truckenbrodt, S., et al., X10 expansion microscopy enables 25-nm resolution on conventional microscopes. EMBO Rep, 2018. 19(9).

4. Damstra, H.G.J., et al., Visualizing cellular and tissue ultrastructure using Ten-fold Robust Expansion Microscopy (TREx). Elife, 2022. 11.

5. Gambarotto, D., V. Hamel, and P. Guichard, Ultrastructure expansion microscopy (U-ExM). Methods Cell Biol, 2021. 161: p. 57–81.

6. Lee, H., et al., Tetra-gel enables superior accuracy in combined super-resolution imaging and expansion microscopy. Sci Rep, 2021. 11(1): p. 16944.

7. Hawkins, T.J., et al., Expansion Microscopy of Plant Cells (PlantExM). Methods Mol Biol, 2023. 2604: p. 127–142.

8. Cho, I. and J.B. Chang, Simultaneous expansion microscopy imaging of proteins and mRNAs via dual-ExM. Sci Rep, 2022. 12(1): p. 3360.

9. White, B.M., et al., Lipid Expansion Microscopy. J Am Chem Soc, 2022. 144(40): p. 18212–18217.

10. Steib, E., et al., TissUExM enables quantitative ultrastructural analysis in whole vertebrate embryos by expansion microscopy. Cell Rep Methods, 2022. 2(10): p. 100311.

11. Gao, M., et al., Expansion Stimulated Emission Depletion Microscopy (ExSTED). ACS Nano, 2018. 12(5): p. 4178–4185.

12. Halpern, A.R., et al., Hybrid Structured Illumination Expansion Microscopy Reveals Microbial Cytoskeleton Organization. ACS Nano, 2017. 11(12): p. 12677–12686.

13. Zwettler, F.U., et al., Molecular resolution imaging by post-labeling expansion single-molecule localization microscopy (Ex-SMLM). Nat Commun, 2020. 11(1): p. 3388.

14. Sheard, T.M.D., et al., Three-Dimensional and Chemical Mapping of Intracellular Signaling Nanodomains in Health and Disease with Enhanced Expansion Microscopy. ACS Nano, 2019. 13(2): p. 2143–2157.

15. Sheard, T.M.D., et al., Three-dimensional visualization of the cardiac ryanodine receptor clusters and the molecular-scale fraying of dyads. Philos Trans R Soc Lond B Biol Sci, 2022. 377(1864): p. 20210316.

16. Shaib, A.H., et al., Expansion microscopy at one nanometer resolution. bioRxiv, 2022: p. 2022.08.03.502284.

17. Karagiannis, E.D., et al., Expansion Microscopy of Lipid Membranes. bioRxiv, 2019: p. 829903.

18. M’Saad, O. and J. Bewersdorf, Light microscopy of proteins in their ultrastructural context. Nature Communications, 2020. 11(1): p. 3850.

19. Sim, J., et al., Nanoscale resolution imaging of the whole mouse embryos and larval zebrafish using expansion microscopy. bioRxiv, 2022: p. 2021.05.18.443629.

20. Konigshausen, E., et al., Imaging of Podocytic Proteins Nephrin, Actin, and Podocin with Expansion Microscopy. J Vis Exp, 2021(170).

21. Gambarotto, D., et al., Imaging cellular ultrastructures using expansion microscopy (U-ExM). Nat Methods, 2019. 16(1): p. 71–74.

22. Pesce, L., et al., Measuring expansion from macro- to nanoscale using NPC as intrinsic reporter. Journal of Biophotonics, 2019. 12(8): p. e201900018.

23. Vanheusden, M., et al., Fluorescence Photobleaching as an Intrinsic Tool to Quantify the 3D Expansion Factor of Biological Samples in Expansion Microscopy. ACS Omega, 2020. 5(12): p. 6792–6799.

24. Damstra, H.G.J., et al., GelMap: Intrinsic calibration and deformation mapping for expansion microscopy. bioRxiv, 2022: p. 2022.12.21.521394.

25. Truckenbrodt, S., et al., A practical guide to optimization in X10 expansion microscopy. Nat Protoc, 2019. 14(3): p. 832–863.

26. Finan, J.D., et al., Nonlinear osmotic properties of the cell nucleus. Ann Biomed Eng, 2009. 37(3): p. 477–91.

27. Zhao, Y., et al., Nanoscale imaging of clinical specimens using pathology-optimized expansion microscopy. Nat Biotechnol, 2017. 35(8): p. 757–764.

28. Suen, K.M., et al., Expansion microscopy reveals subdomains in C. elegans germ granules. Life Sci Alliance, 2023. 6(4).

29. Farnebäck, G. Two-Frame Motion Estimation Based on Polynomial Expansion. in Image Analysis. 2003. Berlin, Heidelberg: Springer Berlin Heidelberg.

